# Spatiotemporal empirical mode decomposition of resting-state fMRI signals: application to global signal regression

**DOI:** 10.1101/556555

**Authors:** Narges Moradi, Mehdy Dousty, Roberto C. Sotero

## Abstract

Resting-state functional connectivity MRI (rs-fcMRI) is a common method for mapping functional brain networks. However, estimation of these networks is affected by the presence of a common global systemic noise, or global signal (GS). Previous studies have shown that the common preprocessing steps of removing the GS may create spurious correlations between brain regions. In this paper, we decompose fMRI signals into 5 spatial and 3 temporal intrinsic mode functions (SIMF and TIMF, respectively) by means of the empirical mode decomposition (EMD), which is an adaptive data-driven method widely used to analyze nonlinear and nonstationary phenomena. For each SIMF, brain connectivity matrices were computed by means of the Pearson correlation between TIMFs of different brain areas. Thus, instead of a single connectivity matrix, we obtained 5 × 3 = 15 functional connectivity matrices. Given the high value obtained for large-scale topological measures such as transitivity, in the low spatial maps (SIMF3, SIMF4, and SIMF5), our results suggest that these maps can be considered as spatial global signal masks. Thus, the spatiotemporal EMD of fMRI signals automatically regressed out the GS, although, interestingly, the removed noisy component was voxel-specific. We compared the performance of our method with the conventional GS regression and to the results when the GS was not removed. While the correlation pattern identified by the other methods suffers from a low level of precision, our approach demonstrated a high level of accuracy in extracting the correct correlation between different brain regions.

## 1. Introduction

Resting-state functional connectivity MRI (rs-fcMRI) has considerable potential for mapping functional brain networks [1, 2, 3, 4, 5, 6]. This mapping, which reveals the brain’s functional architecture and operational principles [1, 2], can be used for early detection of brain connectivity pathologies in neuropsychiatric patients [7]. However, the presence of a common global systemic noise in Blood Oxygen Level Dependent (BOLD) signals fluctuations, known as the global signal (GS), is a significant problem for fcMRI analysis. This presence is problematic as it is of unknown physiological origin [7, 8, 9]. Because of the presence of GS in a functional network, regressing out GS becomes an important step in data preprocessing. Thus, it must be done prior to fcMRI analysis. GS is generally defined as the average of the BOLD signals over the whole brain [9, 10, 11] and can be computed from the raw images or after some preprocessing steps [11]. The average-based GS is called conventional GS (or static GS (SGS)[7]).

Application of SGS regression (SGSR) was at first just limited to task-related fMRI imaging [10, 12]. More recently, SGSR usage has received more attention in the analysis of resting-state fMRI than in task-related fMRI studies [11]. Some studies suggest that application of SGSR improves the functional specificity of resting-state correlation maps and helps to remove non-neuronal sources of global variance like respiration and movement [9, 11, 13]. However, other studies found that these improvements are limited to systems that would exhibit only positive correlations with the specific selected seeds [9, 14]. On the other hand, many studies have shown that the common preprocessing steps of removing GS via a general linear model can create correlations between regions that may never have existed [15, 16, 17, 18], which results in spurious fcMRI values. Moreover, it has been shown that SGSR do not consider the brain’s spatial heterogeneities and biases correlations in different regions of the brain [18]. Accordingly, the extracted correlation maps are known to present artifacts and do not reflect underlying neurological properties [15, 16, 17, 18]. Therefore, regressing out GS is under debate as its removal by applying current approaches may introduce artifacts into the fMRI data or cause the loss of important neuronal components [15, 16, 17, 18].

These concerns about the GSR methods and the need for accurate brain functional connectivity maps motivate the need to develop new methods for dealing with GS. In particular, a method is needed that can reveal accurate relationships between brain regions rather than create spurious correlations and lose physiological information. Here, we define an adaptive global signal regression (AGSR), which is voxel-specific, by performing a spatiotemporal decomposition of the fMRI time series. The Spatial and Temporal Intrinsic Mode Functions (SIMF and TIMF, respectively) of fMRI data are acquired by applying FATEMD [19] and ICEEMDAN [20] methods, respectively. These methods are based on the empirical mode decomposition (EMD) method [21, 22], and will be used to decompose the fMRI signals adaptatively and in a voxel-specific way.

First, the averaged functional connectivity between the extracted peak signals of all brain regions included in the AAL 116 atlas [23] for different TIMFs of each SIMFs over all subjects is computed, obtaining functional connectivity matrices. We then computed the transitivity and efficiency [24, 25, 26] of these matrices. Given the high value of transitivity and efficiency in the low spatial maps (SIMF3, SIMF4, and SIMF5), our results suggest that these maps can be considered as spatial global signal masks. The performance of the proposed method is compared with the SGSR method, and also with the results when GS is not removed. This is done by investigating the functional connections within an extracted peak voxel of the known networks regions and the selected seed region. While the correlation pattern identified by the other methods suffers from a low level of precision, our method demonstrates a high level of accuracy due to its data-driven adaptive nature.

## 2. Methods

### 2.1. fMRI data acquisition

The resting-state fMRI preprocessed data-set of 21 subjects from the NIH Human Connectome Project (HCP) (https://https://db.humanconnectome.org) [27] is used in this research. Each subject was involved in 4 runs of 15 minutes each using a 3 T Siemens, while their eyes were open and had a relaxed fixation on a projected bright cross-hair on a dark background. The data were acquired with 2.0 mm isotropic voxels for 72 slices, TR=0.72 s, TE=33.1 ms, 1200 frames per run, 0.58 ms Echo spacing, and 2290 Hz/Px Bandwidth [28]. Therefore, the fMRI data were acquired with a spatial resolution of 2 × 2 × 2 mm and a temporal resolution of 0.72 s, using multibands accelerated echo-planar imaging to generate a high quality and the most robust fMRI data [28]. The fMRI data were temporally preprocessed and denoised with a highpass filter and ICA-FIX approach [29, 30]. Furthermore, the data were spatially preprocessed to remove spatial artifacts produced by head motion, B0 distortions, and gradient nonlinearities [31]. As comparison of fMRI images across subjects and studies is possible when the images have been transformed from the subject’s native volume space to the MNI space [32, 33], fMRI images were wrapped and aligned into the MNI space with FSL’s FLIRT 12 DOF affine and then a FNIRT nonlinear registration [34, 35, 36]. In this study, the MNI-152-2 mm atlas [37, 38, 39] was utilized for fMRI data registration.

### 2.2. Estimation of the Temporal IMFs (TIMFs)

EMD is an adaptive data-driven signal processing method, which does not need any prior functional basis such as the wavelet transform [22]. The basic functions are derived adaptively from the data by the EMD sifting procedure. The EMD method developed and established by Huang et al. [21] decomposes nonlinear and non-stationary time series into their fundamental oscillatory components, called Intrinsic Mode Functions (IMFs). There are two criteria defining an IMF during the sifting process: 1) the number of extrema and zero crossings must be either equal or differ at most by one, and, 2) at any instant in time, the mean value of the envelope defined by the local maximum and the envelope of the local minimum is zero. The EMD algorithm for estimating the IMFs of the time series *x*(*t*) is:

1. *r*_0_(*t*) = *x*(*t*), *j* = 1.
2. For extracting the *j*-th IMF:

a. *h*_0_(*t*) = *r*_*j*_(*t*), *k* = 1,
b. Locate local maximum and minimum of *h*_*k*−1_(*t*),
c. Identify the averaged envelope using cubic spline interpolation to define upper and lower envelope of *h*_*k*−1_(*t*),
d. Calculate the mean value *m*_*k*−1_(*t*),
e. Put *h*_*k*_(*t*) = *h*_*k*−1_(*t*) − *m*_*k*−1_(*t*),
f. Check the stopping criteria. The stopping criteria determines the number of sifting steps to decompose an IMF [21]. If stopping criteria is satisfied then *h*_*j*_(*t*) = *h*_*k*_(*t*) otherwise, go to (a) to extract next IMF with *k* = *k* + 1.
3. *r*_*j*_(*t*) = *r*_*j*−1_(*t*) − *h*_*j*_(*t*).
4. If at least two extrema were in the *r*_*j*_(*t*), the next IMF is extracted otherwise the EMD algorithm is finished and *r*_*j*_(*t*) is the residue of *x*(*t*). Accordingly, *x*(*t*) is defined as:

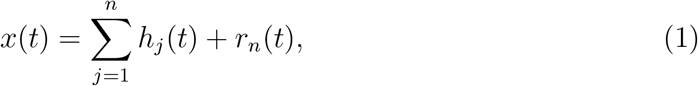

where *h*_*j*_(*t*) is the *j*-th IMF, *n* is the number of IMFs, and *r_n_*(*t*) is the residue of the signal. Thus, the EMD method adaptively decomposes a time series into a set of IMFs and a residue where the first IMF (IMF1) corresponds to the fastest oscillatory mode and the last IMF (IMFn) to the slowest one [21, 40]. However, frequent occurrences of the mode-mixing phenomenon in analyzing real signals using EMD algorithm is problematic. To address this problem and improve the spectral separation of modes, the ensemble empirical mode decomposition (EEMD) method was proposed [41]. This method extracts modes by performing the decomposition over an ensemble of noisy copies of the original signal combined with white Gaussian noises, and taking the average of all IMFs in the ensemble [20].

The EEMD method solves the mode mixing problem, but certain issues remain. First, the number of IMFs extracted from each of the noisy signal copies is different, and this creates a problem when averaging the IMFs. The second problem is a reconstruction error in the EEMD method[20, 41]. To fix this error the complementary EEMD (CEEMD) was proposed [42]. In the CEEMD algorithm, pairs of positive and negative white noise processes are added to the original signal to make two sets of ensemble IMFs. Accordingly, the CEEMD effectively eliminates residual noise in the IMFs which alleviate the reconstruction problem. Nonetheless, the problem of the different number of modes when averaging still persists. To overcome this problem, the CEEMD with adaptive noise (CEEMDAN) was developed [20, 43]. In this approach, the first mode is computed exactly as in EEMD. Then, for the next modes, IMFs are computed by estimating the local means of the residual signal plus different modes extracted from the white noise realizations. CEEMDAN decomposition can create some spurious modes with high-frequency and low-amplitude due to overlapping in the scales. Additionally, some residual noise is still present in the modes. As a consequence, the new optimization algorithm, Improved Complete Ensemble Empirical Mode Decomposition with Adaptive Noise (ICEEMDAN), was proposed [20].

During the sifting process using ICEEMDAN method the local mean of realizations is estimated, instead of using the average of modes from the first step. This change in the algorithm reduces the amount of noise present in the final computed modes. To deal with the issue of creation of spurious modes in the final results, ICEEMDAN method proceeds differently than the EEMD and CEEMDAN methods. In ICEEMDAN, white noise is not added directly; instead EMD modes of white noise are added to the original signal and to the IMFs during the sifting process [20, 41]. Furthermore, in this method as in CEEMDAN, a constant coefficient is added to the noise that makes the desired signal to noise ratio between the added noise and the residue to which the noise is added. This coefficient is computed based on the standard deviation of the residue at each step of the sifting process. Therefore, the IMFs computed with ICEEMDAN have less noise and more physical content than IMFs obtained with other methods [20] (More detailed description of ICEEMDAN method can be found at [20]). The high accuracy rate, reduction in the amount of noise contained in the modes, and the alleviation of mode mixing phenomenon qualify this method to effectively decompose biological signals. In this paper the ICEEMDAN method with 300 ensembles and a level of noise of 0.2 is used to extract the Temporal Intrinsic Mode Functions (TIMFs) from the fMRI data.

### 2.3. Estimation of the Spatial IMFs (SIMFs)

A fast, time efficient, and effective method is essential for processing real images that have a large size. Previous EMD-based methods were limited to small size images as the extrema detection, interpolation at each iteration, and the large number of iterations make their processing time consuming and complicated [19, 44, 45, 46]. Therefore, those methods were just applicable to reduced size images, which resulted in losing some information during their process. Fast and Adaptive Tridimensional (3D) EMD, abbreviated as FATEMD, is a recent extension of the EMD method to three dimensions [19]. The FATEMD method is able to estimate volume components called tridimensional Intrinsic Mode Functions (3D-IMFs) quickly and accurately by limiting the number of iterations per 3D-IMF to one, and changing the process of computing upper and lower envelopes, which reduce the computation time for each iteration [19, 44, 46]. In the FATEMD method, the steps of extracting 3D-IMFs are almost the same as the previous EMD based methods, except for the number of iterations and the estimations of the maximum and minimum envelopes. The steps for decomposing a volume *V* (*m, n, p*) with dimensions *m, n*, and *p* using the FATEMD approach are as follows [19, 44]:

1. Set *i* = 1, *R*_*i*_(*m, n, p*) = *V* (*m, n, p*).
2. Determine the local maximum and minimum values by browsing *R*_*i*_(*m, n, p*) using a 3D window (cube) with a size of 3 × 3 × 3 which results in an optimum extrema maps (Map_max_(*m, n, p*) and Map_min_(*m, n, p*)). These local maximum (or minimum) values are strictly higher (or lower) than all of their neighborhoods contained in the cube.
3. Calculate the size of the Max and the Min filters which will be used in making extrema envelopes and their smoothness. The maximum and minimum filters are made by computing the nearest Euclidean distances between the maximum (*d*_adj.max_) (minimum (*d*_adj.min_)) points. The cubic window width (*w*_en_) then is determined by using one of the following four formulae for both maximum and minimum filters. Here, we used the 4-th formula as outlined below, although using the other formulas will result in approximately the same decomposition result:

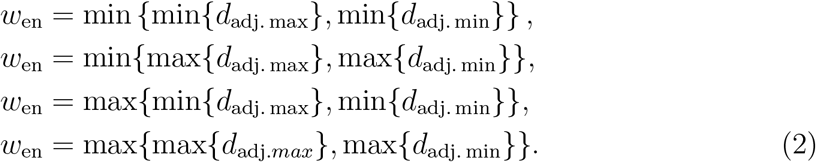
4. Create the envelopes of maxima and minima (Env_max_(*m, n, p*) and Env_min_(*m, n, p*)) of size (*w*_en_).
5. Use the mean filter to compute the smoothed envelopes: Env_max−s_(*m, n, p*) and Env_min−s_(*m, n, p*).
6. Calculate the mean filter by averaging the smoothed upper and lower envelopes(Env_*A*_(*m, n, p*)).
7. Calculate the *i*^*th*^ 3D-IMF : IMF_*i*_(*m, n, p*) = R_*i*_(*m, n, p*) − Env_*A*_(*m, n, p*).
8. Calculate R_*i*+1_(*m, n, p*) = R_*i*_(*m, n, p*) − IMF_*i*_(*m, n, p*).
9. If R_*i*+1_(*m, n, p*) contains more than two extrema then

Go to the step 2 and set *i* = *i* + 1, Else

The FATEMD decomposition is completed.

Therefore, FATEMD is an adaptive approach as all of the processes for computing filters and making the maximum, minimum, and the mean envelops are data driven. FATEMD decomposes a volume into a set of 3D-IMFs [19]. In general, a volume V can be reconstructed from the K 3D-IMFs and the residue as follows:

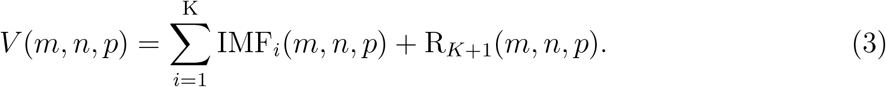

*K* is the number of IMFs, and *R*(*m, n, p*) is the residue of the signal.

In this paper, we apply the FATEMD method at each time instant to decompose the resting-state fMRI data into tridimensional IMFs called Spatial Intrinsic Mode Functions (SIMF). Fig. (1) shows the spatial decomposition results of a sample resting-state fMRI image. The ICEEMDAN method is then utilized to decompose each SIMF into its corresponding TIMFs.

**Figure 1:**
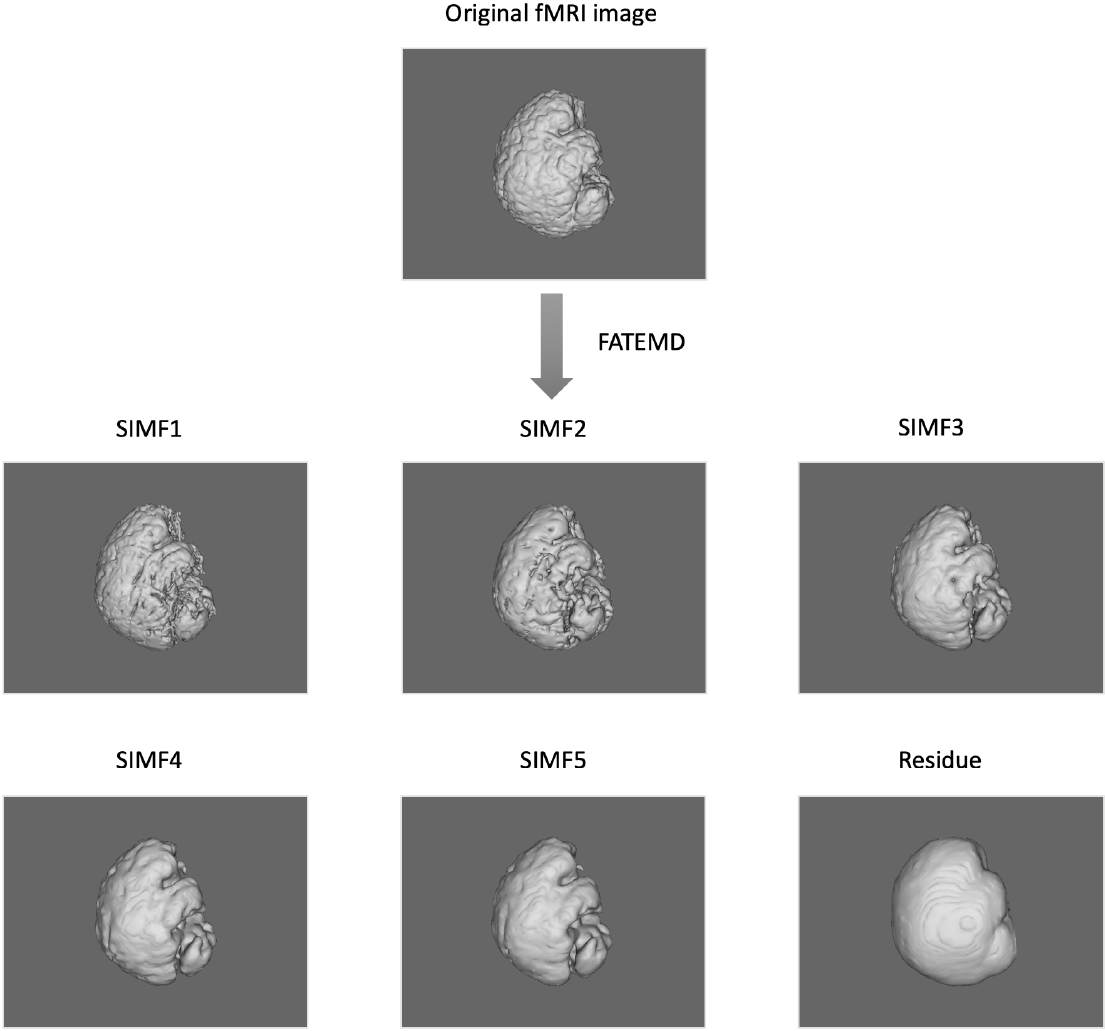
Spatial decomposition of a sample fMRI image using FATEMD method. The original fMRI image at one TR time is decomposed into 5 SIMFs (SIMF1 to SIMF5) and a residue.

### 2.4. Spatiotemporal pattern analysis of the fMRI data

To define an adaptive and voxel-specific GS, we investigate the spectral information of fMRI data by constructing the functional connectivity matrices using extracted TIMFs and SIMFs data. To fulfill this aim, first, the SIMFs of the fMRI data at each TR time are computed by applying the FATEMD method, then, all spatial components are merged together in time to construct the time series of each SIMF. Second, the peak voxel in each region, that is, the voxel of maximal activation, is selected since it provides the best effect of any voxel in the ROI [47]. Additionally, the peak voxel activity correlates better with evoked scalp electrical potentials than approaches that average activity across the ROI. This means that the peak voxel represents the ROIs activity better than other choices [48]. The peak voxel in each region is determined using previously published Talairach coordinates (after conversion to MNI coordinates and using AAL 116 atlas) [49]. After determining the peak voxels of each region, the ICEEMDAN method is applied to its time series to compute the TIMFs. Thus, the TIMFs of all regions for each SIMF are computed.

We then compare the predefined distinct frequency bands presented in fMRI studies (slow5 [0.01-0.027 Hz], slow4 [0.027-0.073 Hz], slow3 [0.073-0.198 Hz], slow2 [0.198-0.25 Hz], and slow1 [0.5-0.75 Hz]) [50, 51], to the frequency content of the extracted TIMFs. As seen in the Fig. (2), the frequency range comprised in TIMF1 to TIMF3 is approximately the same as the frequency range of the combination of slow1 to slow3. The frequency range of TIMF4 is the same as slow4, and the frequency range of the combination of TIMF5 to TIMF9 has the same frequency range as the slow5 frequency band. Accordingly, we label the combination of TIMF1 to TIMF3 as TIMF1, TIMF4 as TIMF2, and the combination of TIMF5 to TIMF9 as TIMF3. Fig. (3) represents the pipeline used in computing SIMF and TIMF for each resting-state fMRI data. Accordingly, the functional connectivity matrices are constructed by computing the average of correlation coefficients between all possible pairs of TIMFs correspond to different Spatial domains for all brain regions comprised in the AAL 116 atlas over all 21 subjects. Consequently, instead of the classical functional connectivity matrix, the decomposition presented here produces 5 × 3 = 15 connectivity matrices (each with size 116 × 116), 3 temporal domains and 5 spatial domains, encompassing the rich spatiotemporal dynamics of brain activity.

**Figure 2:**
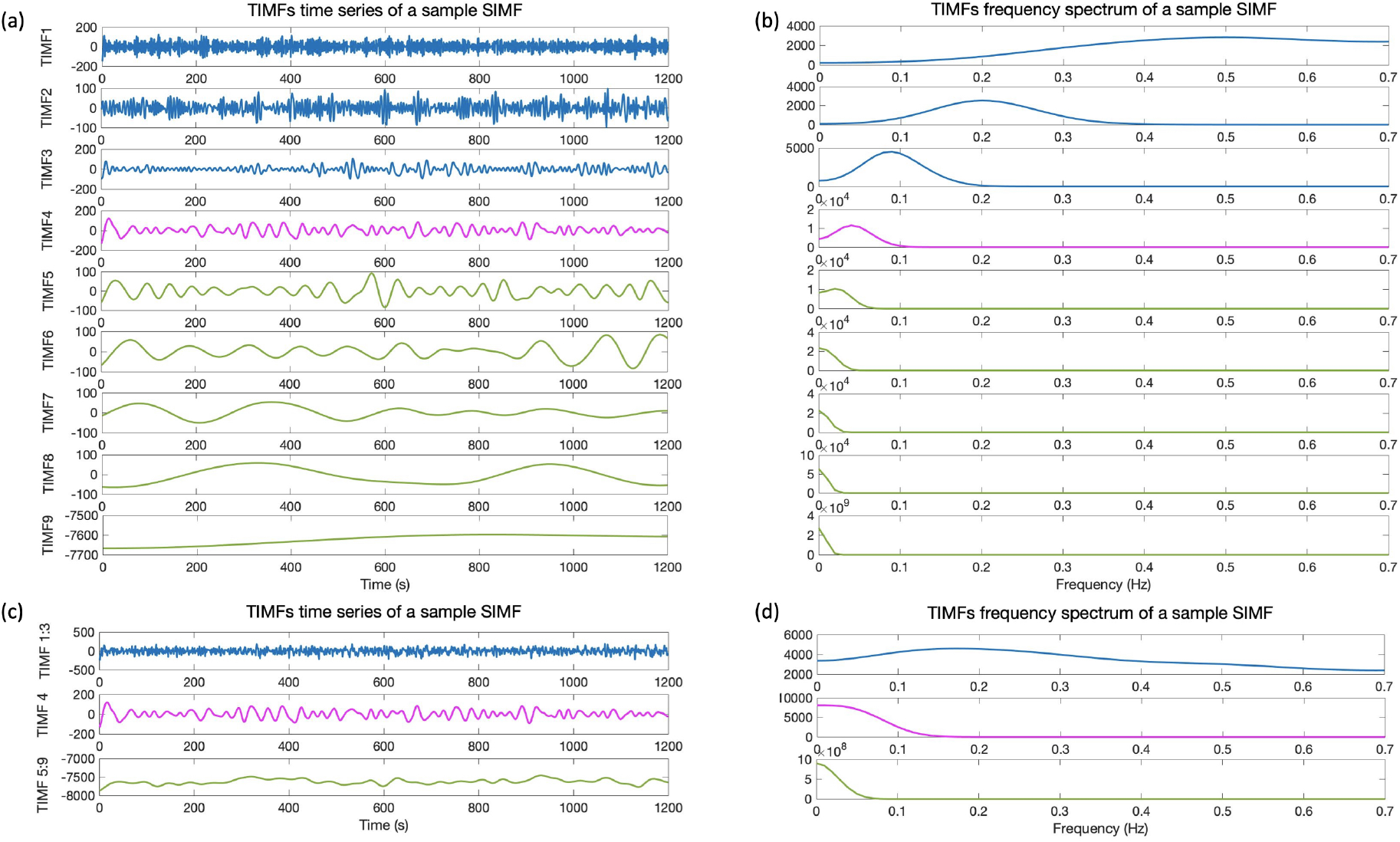
Temporal IMFs and their corresponding frequency spectrum. a) 9 decomposed TIMFs of a sample SIMF by applying the ICEEMDAN method with 300 ensembles and a level of noise of 0.2. b) Represents the frequency spectrum of the 9 TIMFs. c) The 9 decomposed TIMFs are divided into three different frequency bands. According to slow1 to slow3 and slow5 frequency bands defined in the literature, TIMFs1 to 3 and 5 to 9 are combined, respectively. d) The frequency spectrum of TIMFs in part (c) that correspond to frequency bands used in the literature for slow1 to slow5 [50, 51]. TIMF : Temporal Intrinsic Mode Function, ICEEMDAN: Improved Complete Ensemble Empirical Mode Decomposition with Adaptive Noise.

**Figure 3:**
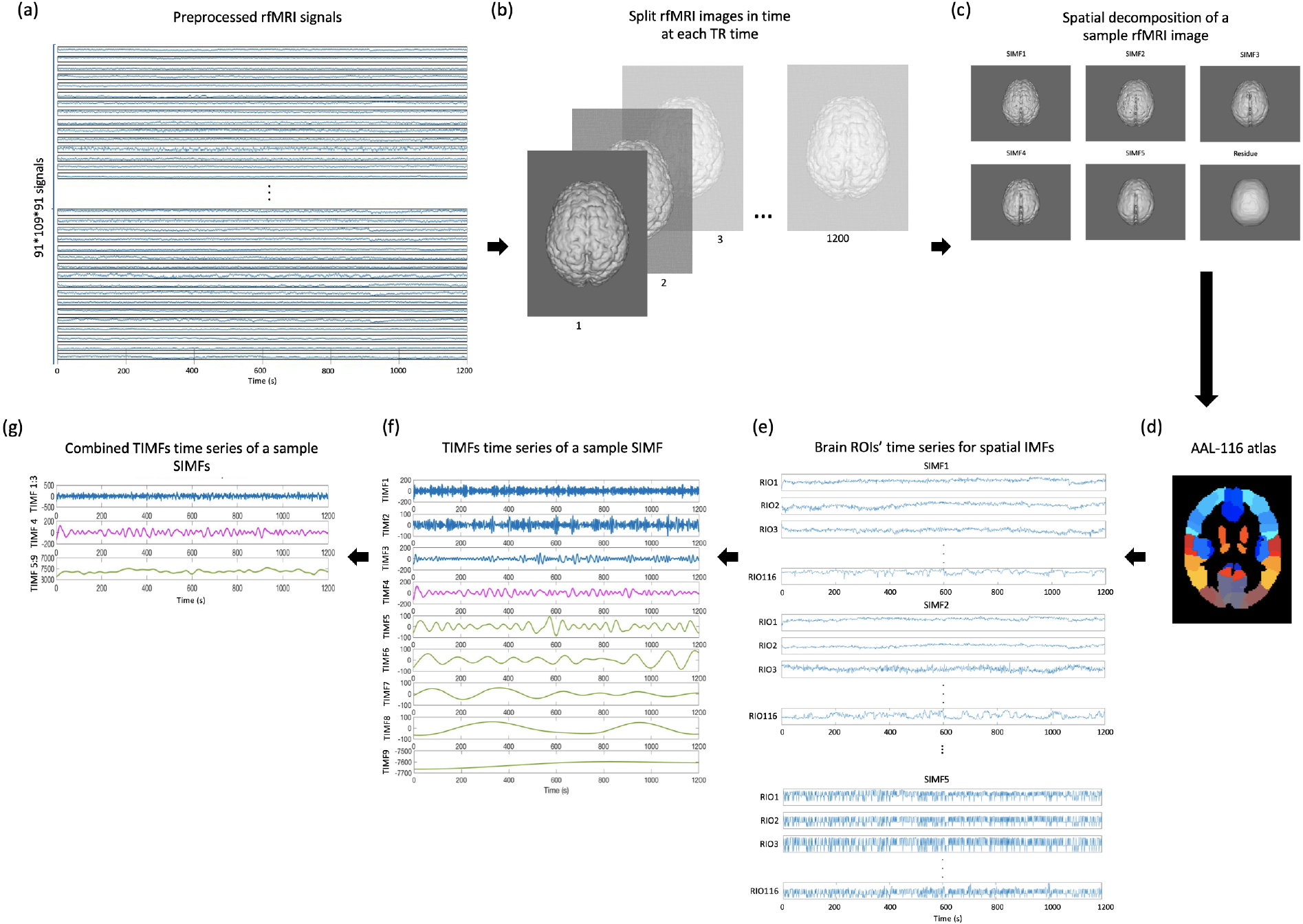
Pipeline for computing spatial and temporal IMFs (SIMF and TIMF) of the fMRI data. a) A sample of fMRI data. b) Splitting each fMRI data in time at each TR time. c) SIMFs at each TR time which are computed by applying FATEMD approach. d) Shows the AAL 116 atlas used after merging SIMFs in time to select the peak voxel of each region. e) Time series of all brain ROIs for each SIMF. f) The TIMFs’ time series of a sample SIMF for one ROI computed by using ICEEMDAN approach. g) Combination of time series of the TIMFs in (f) based on frequency bands of slow1 to slow5 defined in the literature. The combination of the TIMF1 to TIMF3, TIMF4, and the combination of TIMFs5 to TIMF9 are labeled as TIMF1, TIMF2, and TIMF3 in the rest of the paper, respectively. rfMRI: resting-state fMRI, TIMF: Temporal Intrinsic Mode Function, SIMF: Spatial Intrinsic Mode Function, ICEEMDAN: Improved Complete Ensemble Empirical Mode Decomposition with Adaptive Noise, FATEMD: Fast and Adaptive Empirical Mode Decomposition, ROI: Region of Interest.

### 2.5. Integration and segregation of functional connectivity matrices

In studying brain networks, the efficiency of the information exchange in the network is examined at two scales, global and local. Generally, integration is based on the concept of a path between regions that estimates the simplicity of the brain regions’ communication [24, 25]. The efficiency between two nodes is defined by computing the shortest path length. The shortest path length is computed by counting the smallest number of edges needed to get from node *i* to node *j*. The shortest path length is inversely related to node weight, as strong association has a large weight which shows a shorter length and close proximity. Functional connectivity matrices provide the information needed to estimate the weight and the shortest path length between all pairs of brain regions [24, 26].

The global efficiency, a measure of network’s integration, is computed by taking the average inverse of the efficiency of all the node pairs of the network that is normalized by the maximal number of network’s links. Therefore, the weighted global efficiency measures how efficient the brain is able to combine information from different brain regions and is computed via the following equation:

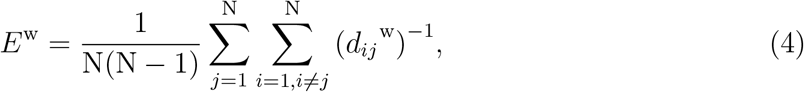

where N is the number of nodes in the network and *d*_*ij*_is the minimum path length between nodes *i* and *j*. When two nodes are disconnected the length of that path would be infinite and correspondingly, the efficiency would be zero [24, 25]. On the other hand, the local efficiency measures the averaged efficiency of information transfer in the neighboring nodes of the node *i* in network excluding node *i* itself. It is associated with functional segregation of the network and characterization the pattern of local anatomical circuitry [25, 26]. In other words, local efficiency shows the presence of clusters and modules (interconnected groups) within the network. Clustering coefficient of the network is defined as the fraction of triangles around each node. The mean clustering coefficient for the network shows the clustered connectivity around each node of the network. The normalized and classical variant of the clustering coefficient known as transitivity is an important source of high clustering coefficient in real-world networks which is defined for individual node and describe the presence of densely interconnected groups of regions [52]. The weighted transitivity is computed as:

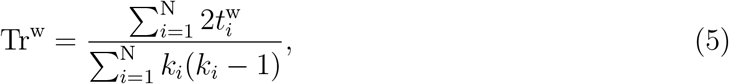

where the *t*_*i*_ is the number of triangles around node *i* and *k*_*i*_ is the degree of the node *i* [24, 26]. We compute the integration and segregation of functional connectivity matrices to assess the power of communication and the density of interconnected groups of brain regions under different conditions.

## 3. Results

### 3.1. Defining Adaptive Global Signal (AGS)

We computed the functional connectivity matrices between all pairs of brain regions for different spatiotemporal domains extracted from fMRI data for each subject. Fig. (4) shows the average connectivity matrix over the 21 subjects. As seen in the figure, SIMF1 and SIMF2 in all TIMFs showed low connectivity whereas SIMF3 to SIMF5 in all TIMFs showed high connectivity. Besides, it indicates that the magnitude of the correlation does not significantly depend on the TIMFs. Thus, based on the connectivity strength for different spatiotemporal domains, the combination of the SIMF1 to SIMF2 and the SIMF3 to SIMF5 including all TIMFs, were considered as two separate signals. We also averaged the six connectivity matrices resulting from the combination of TIMF1 to TIMF3 with SMF1 and SMF2 (Fig. (4)) and labelled it as AGSR (Fig. (5a)), and the nine connectivity matrices resulting when combining TIMF1 to TIMF3 with SIMF3 to SIMF5, which we labelled as AGS (Fig. (5b)). Finally, we computed the average connectivity matrix over resting-state fMRI data of all subjects when no GSR was performed (Fig. (5c)) and labelled it as NR (No Regression).

**Figure 4:**
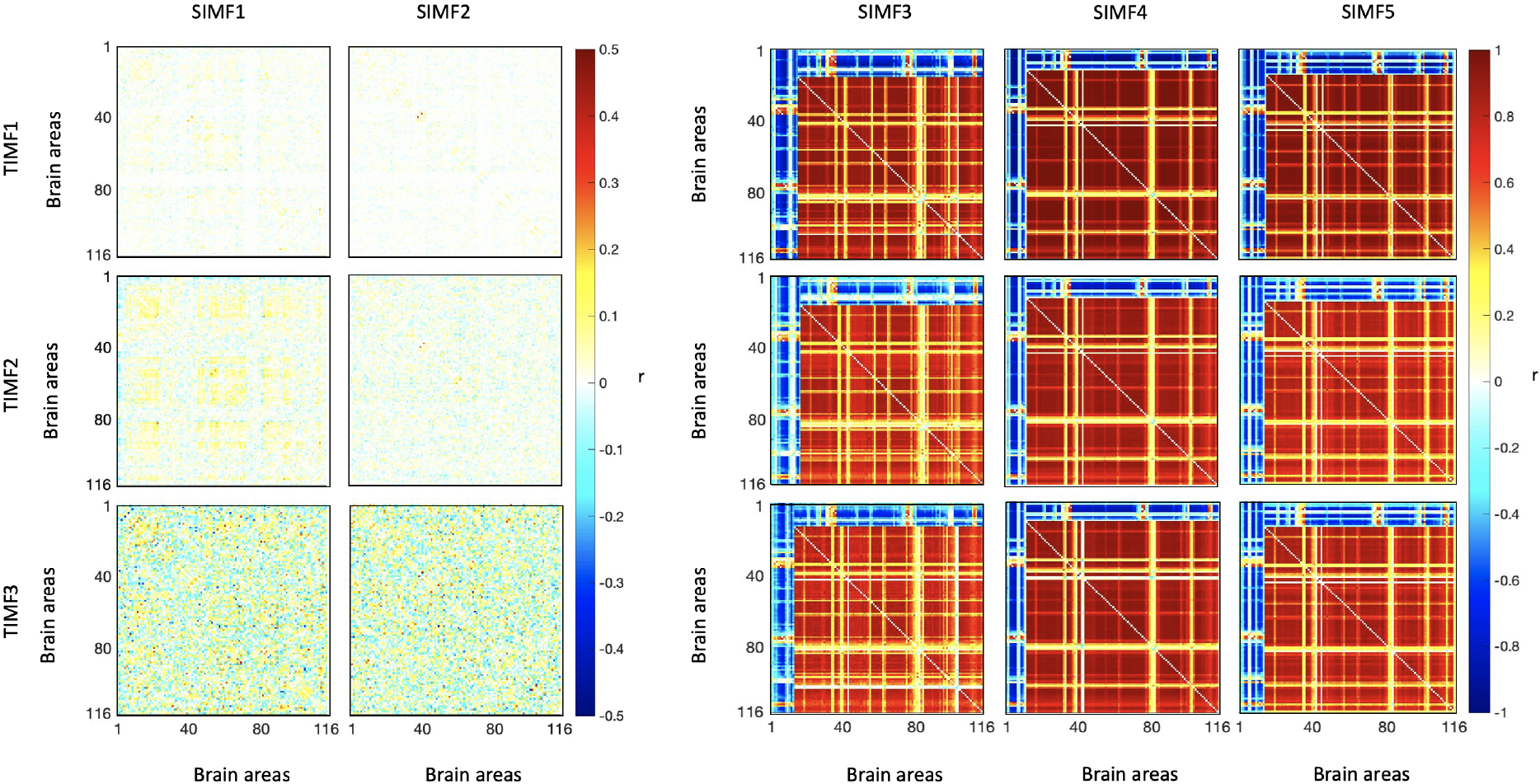
Functional connectivity matrices of the whole brain regions using AAL 116 atlas for different spatial and temporal IMFs. Pearson’s correlation coefficient (*r*) with *P* ≤ 0.01 is computed between all the brain regions’ spatiotemporal domains extracted from fMRI data. Spatial domains are extracted by applying FATEMD method on fMRI signal. The three temporal domains including TIMF1, TIMF2, and TIMF3 are computed by applying ICEEMDAN on each SIMF. SIMF: Spatial Intrinsic Mode Function, TIMF: Temporal Intrinsic Mode Function, ICEEMDAN: Improved Complete Ensemble Empirical Mode Decomposition with Adaptive Noise, FATEMD: Fast and Adaptive Empirical Mode Decomposition.

**Figure 5:**
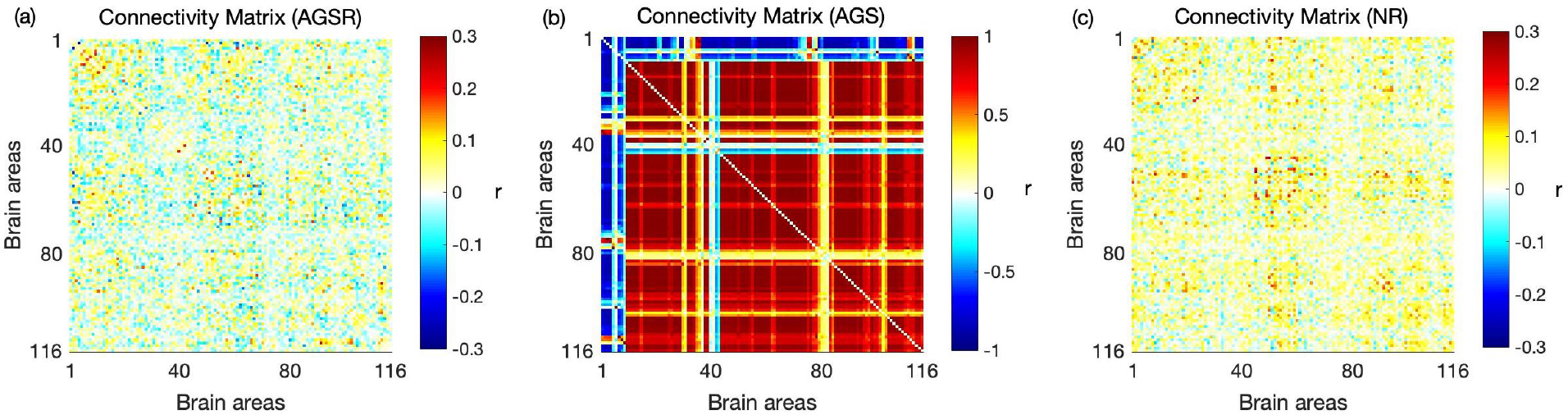
Averaged functional connectivity matrices of the whole brain regions using AAL 116 atlas over all subjects. a) Averaged connectivity matrix of fMRI data when NR is done, b) Averaged connectivity matrix of fMRI data applying AGSR which means the connectivity matrix of combination of SIMF1 and SIMF2 with all TIMFs of the fMRI data, and c) connectivity matrix of the AGS which is the combination of SIMF3 to SIMF5 with all TIMFs. NR: No Regression, AGSR: Adaptive Global Signal regression, AGS: Adaptive Global Signal, SIMF: Spatial Intrinsic Mode Function, TIMF: Temporal Intrinsic Mode Function.

We also computed the transitivity and efficiency for different spatial and temporal IMFs using Eq. (4) and Eq. (5) and also based on functional connectivity results (Figs. (6a), (6b)). Figs. (6a), (6b) show that there are high level of transitivity, and efficiency in the low frequencies of spatial domains, SIMF3, SIMF4, and SIMF5, which indicate active shared connections between all the nodes in the brain, suggesting the existence of GS in the low-frequency spatial domain, called Adaptive Global Signal(AGS).

As seen in Figs. (6c), (6d), and Table (1) the magnitude of averaged transitivity and efficiency of the combination of SIMF1 and SIMF2 with TIMF1 to TIMF3 are almost the same as when no GS is removed from the fMRI time series.

**Figure 6:**
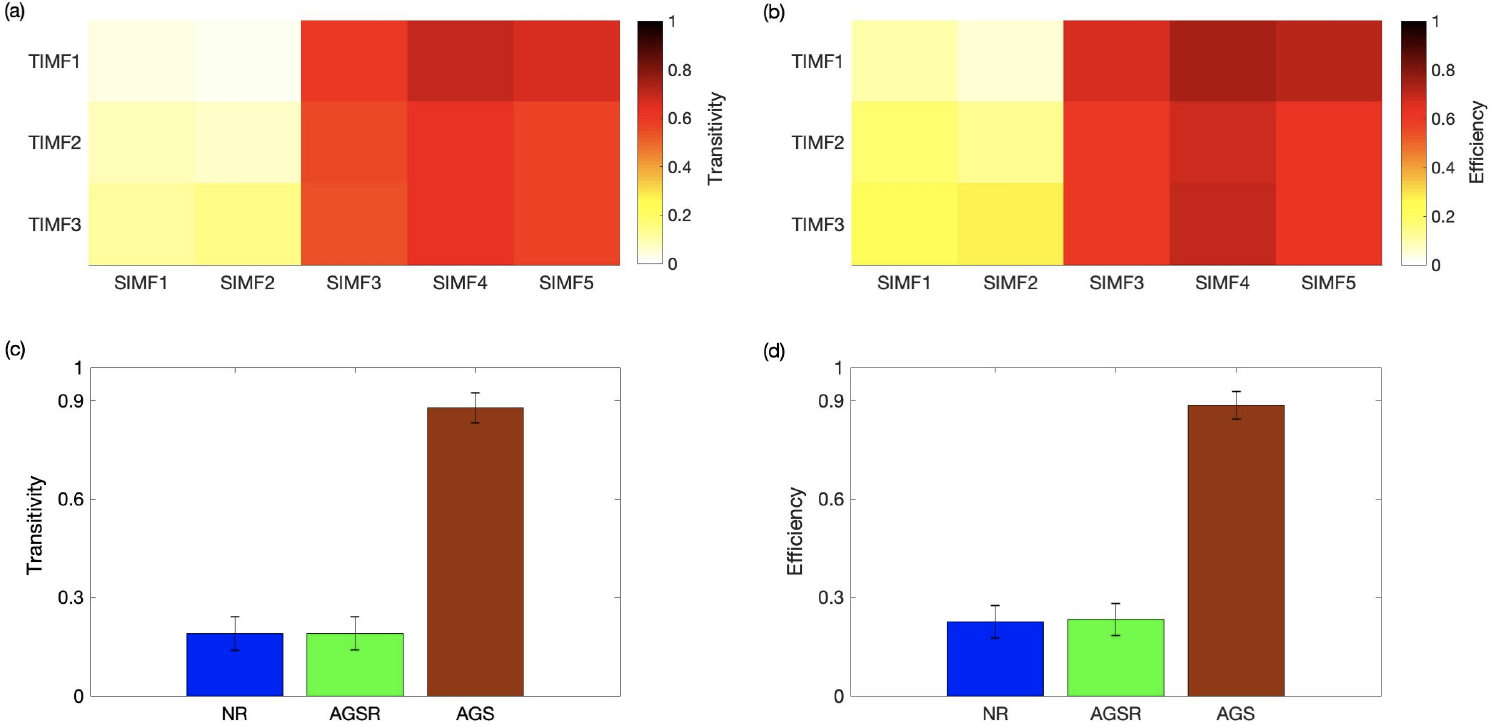
Map of transitivity and efficiency. a) Transitivity and b)Efficiency of the whole brain network for different spatial and temporal IMFs defined in functional connectivity. Comparing the magnitude of c) averaged transitivity and d) averaged efficiency of the brain network over all subjects when the GS from the resting-state fMRI data is not removed, when the AGS is removed (combination of SIMF3 to SIMF5 in all TIMFs is removed from the fMRI signal), and the averaged transitivity and efficiency of the AGS (combination of SIMF3 to SIMF5 in all TIMFs). High values for transitivity and efficiency in the SIMF3 to SIMF5 which represent the AGS in the fMRI data are seen in the figures. NR: No Regression, GS: Global Signal, AGS: Adaptive GS.

**Table 1:** Integration and segregation of functional connectivity matrices under different conditions (NR, AGSR, and AGS). The averaged transitivity and efficiency of the brain network over all subjects, when the NR and the AGSR are performed, and the averaged transitivity and efficiency of the AGS. NR: No Regression, AGS: Adaptive Global Signal, AGSR: AGS Regression.

| Network Measure | Label | Interpretation | Value |
| --- | --- | --- | --- |
| Transitivity | NR | The brain network's averaged transitivity when NR is performed | $0.1903 \pm 0.0513$ |
| Transitivity | AGSR | The brain network's averaged transitivity when AGSR is performed | $0.1902 \pm 0.0510$ |
| Transitivity | AGS | The brain network's averaged transitivity of the AGS | $0.8766 \pm 0.0461$ |
| Efficiency | NR | The brain network's averaged efficiency when NR is performed | $0.2252 \pm 0.0496$ |
| Efficiency | AGSR | The brain network's averaged efficiency when AGSR is performed | $0.2325 \pm 0.0480$ |
| Efficiency | AGS | The brain network's averaged efficiency of the AGS | $0.8850 \pm 0.0417$ |

This implies that the low frequency spatial domains do not play a significant role in the magnitude of integration and segregation of the brain’s networks and just cause spurious connectivity results between brain regions. Furthermore, a high level of integration and segregation in these domains (Figs. (6c), (6d), and Fig. (5b)) confirm that they can be considered as a GS which has to be removed from the fMRI data to have accurate results.

### 3.2. Regressing out the AGS and SGS from fMRI data

According to the definition of AGS, for each brain voxel signal, there is a corresponding AGS while the SGS is common for the whole brain voxels. The AGS for each voxel is computed by summing up the SIMF3, SIMF4, and SIMF5 with all TIMFs while the SGS is computed by taking the average of all brain voxels’ time series. It should be noted that in computing AGS, the residues of spatiotemporal decomposition of the fMRI data are added to last TIMF and SIMF. The three time courses in Figs. (7a), (7b), and (7c) correspond to the AGS, the fMRI sample time course (the peak voxel’s time course in Lateral Parietal cortex (LP) ROI), and the conventional or Static GS (SGS), respectively. Figs. (7d) and (7e) show resting-state fluctuations of the sample fMRI time series from LP ROI after regressing out the AGS and the SGS.

**Figure 7:**
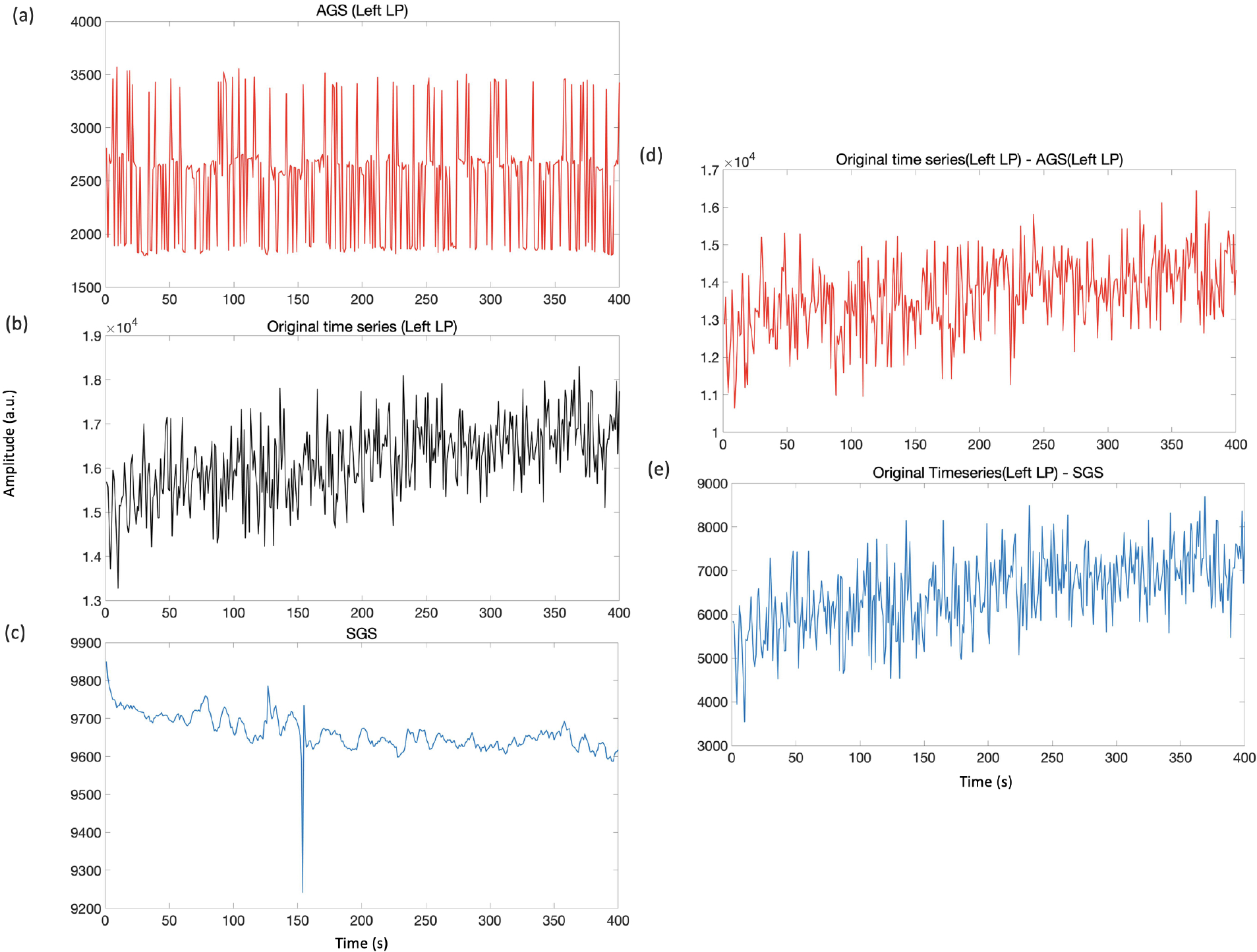
AGSR and SGSR of a sample fMRI data. a) The voxel-specific AGS and b) the original fMRI time series of the peak voxel in lateral parietal(LP) cortex region. c) The SGS which is common for all region’s voxels. d, e) show the time series with the SGSR and AGSR in a time window of 400 s, respectively. These time series are computed by subtracting the AGS and SGS from the original time series. The first 400 time points are shown in the figure for illustration purposes. LP: Lateral Parietal cortex, AGS: Adaptive Global Signal, SGS: Static(conventional) Global Signal, AGSR: AGS Regression, SGSR: SGS Regression.

### 3.3. Connectivity map of task-positive and task-negative networks

The default mode network is a state of brain activation whereby the individual is not attending to any external cues in the environment but certain brain regions are still activated and they are less active during task performance rather than during the resting-state. It has been shown that [49] the default network responses are significantly activated in three of the seeded regions: the Posterior Cingulate Cortex (PCC), Medial Prefrontal cortex (MPF), and Lateral Parietal cortex (LP). The efficacy of our approach is examined by computing the connectivity map. We computed the averaged connectivity between the time course of the PCC region as a seed region and the main regions of the Task Positive Network (TPN) which are the Middle Temporal (MT), right Frontal Eye Field (FEF), left Intraparietal Sulcus (IPS), Visual regions, and the left Auditory region and the Task Negative Network (TNN) ROIs which are MPF, PCC, and left LP which includes the Angular Gyrus, Hippocampus, and Cerebellar tonsils ROIs [7, 9].

Considering the AGS definition, the summation of the SIMF1 and SIMF2 was used to compute the functional connectivity between PCC and TNN and TPN including visual ROIs by using the Pearson’s correlation coefficient (r), *P* ≤ 0.01. Fig. (8) is functional connectivity brain map for different brain layers along the Z axis which show the mean connectivity over all subjects between brain regions and the PCC ROI as a seed region when the AGSR, NR, and the SGSR are performed.

Functional connectivity between different ROIs in Fig. (9), show correct averaged connectivity between the PCC ROI and different regions of the TPN and the TNN applying the new approach of GS in resting-state fMRI data.

While the NR and SGSR (conventional GSR which is based on averaging) are unable to identify the accurate connectivity in some regions for TPN and TNN ROIs, the AGSR approach obtains correct functional connectivity for all regions in TNN and TPN which confirms the effectiveness of the proposed method for GSR (Fig. (9)). As AGSR is an adaptive and voxel-specific method, we have a unique local signal for each voxel which by being removed from fMRI data augments the precision of the rsfc-MRI results.

**Figure 8:**
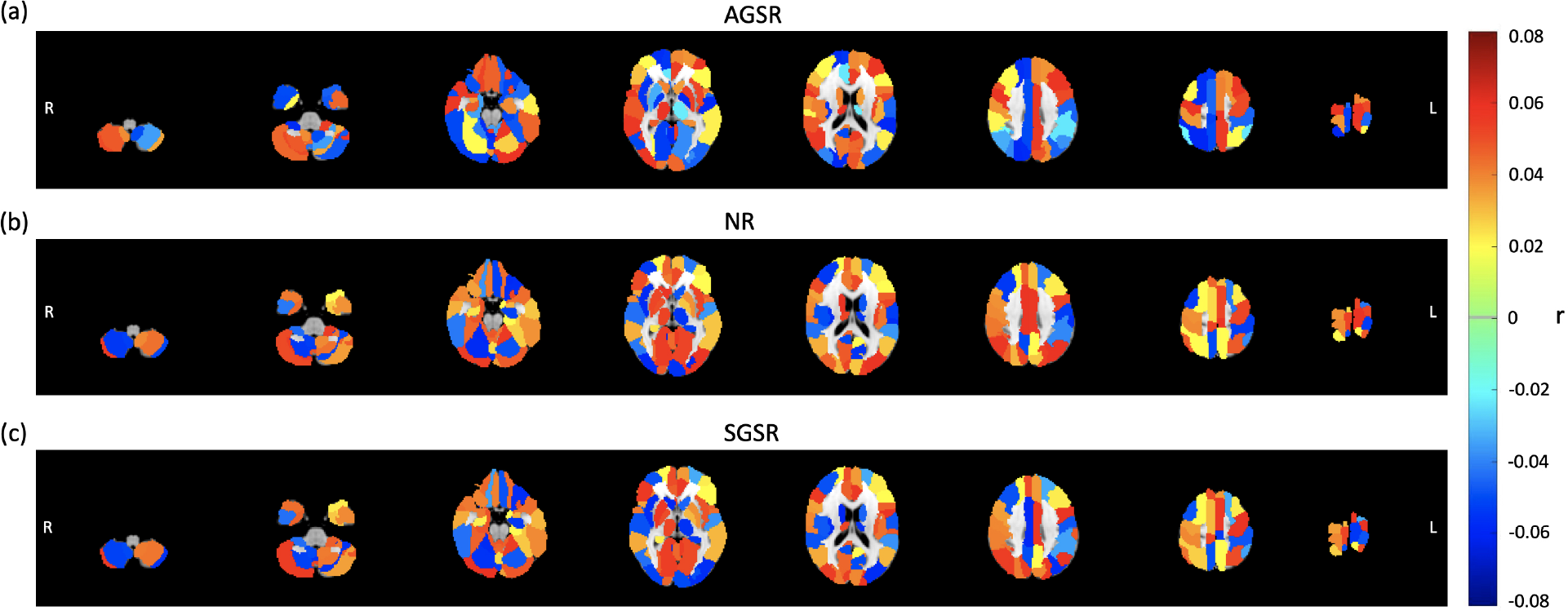
Comparing the averaged functional connectivity between the PCC ROI as a seed region and the brain ROIs using the AAL 116 atlas for fMRI data of all subjects. The averaged functional connectivity applying a) AGSR, b) NR, and c) SGSR. Slices shown in the maps are at Z = 09, 15, 25, 35, 45, 55, 65, 75, respectively. AGSR: Adaptive Global Signal Regression, NR: No Regression, SGSR: Static (conventional) Global signal regression.

**Figure 9:**
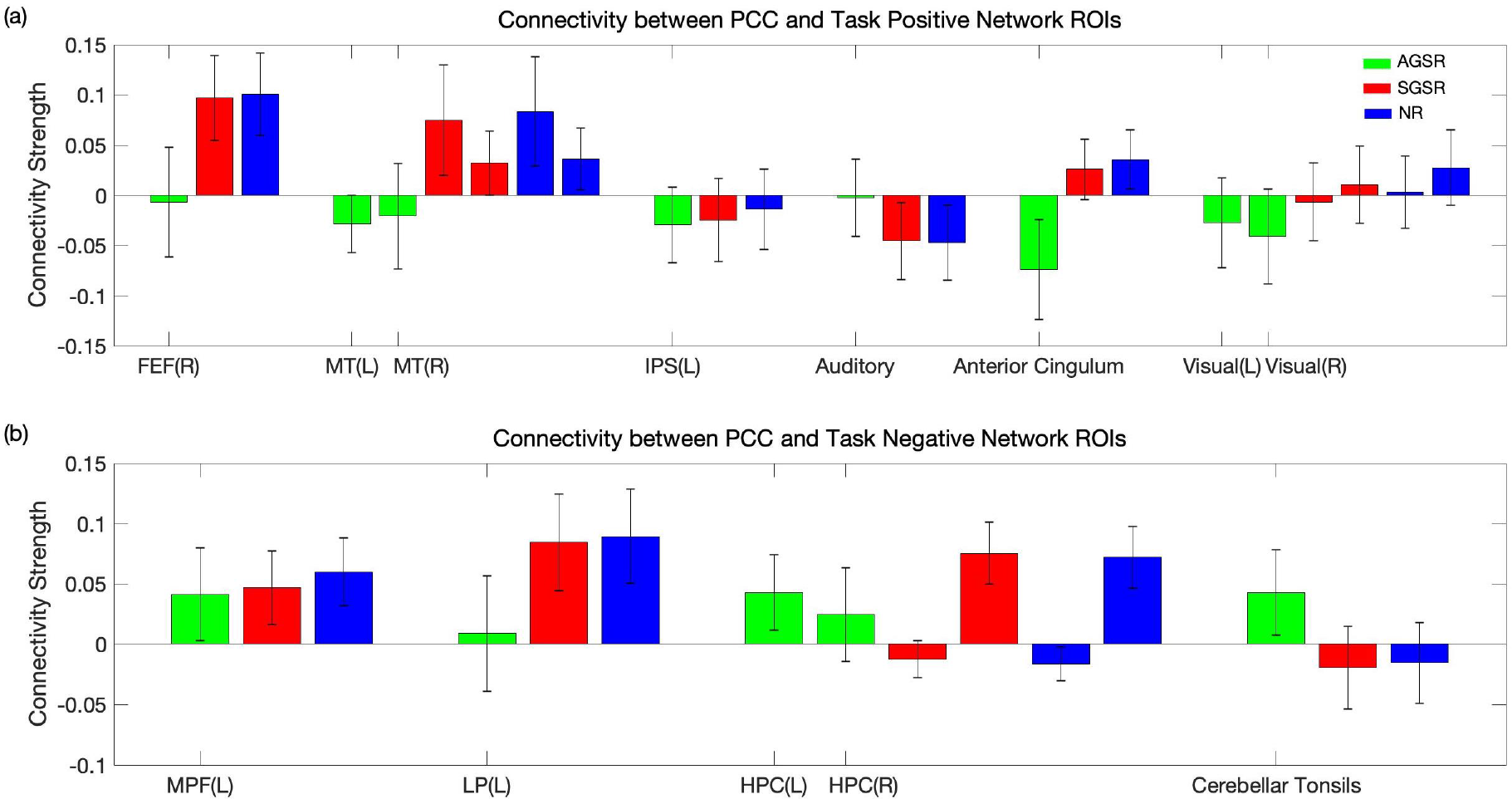
The averaged connectivity map between the PCC as a seed region and a) the TPN and b) TNN ROIs for fMRI data of all subjects. Connectivity results of applying AGSR and SGSR are shown in green and red, respectively, and the blue ones are the results of computing connectivity without applying any GSR (NR). Connectivity map is made by computing Pearson’s correlation coefficient (r) with *P* ≤ 0.01 between the PCC region as a seed region and the main regions of the TPN and TNN. PCC: Posterior Cingulate Cortex, MPF: Medial Prefrontal cortex, LP: Lateral Parietal cortex, MT: Middle Temporal, FEF: Frontal Eye Field, IPS: Intraparietal Sulcus, ROI: Region Of Interest, NR: No Regression, AGS: Adaptive Global Signal Regression, SGS: Static (conventional) Global Signal Regression.

## 4. Discussion

In contrast to previous works [9, 10, 11], the present study provides a new method for GSR, called AGSR, that works voxel-specifically and adaptively. It is believed that fMRI data are a superposition of the GS and network-specific fluctuations. However, the main reason for the controversy over the use of GS in fMRI studies is that the average-based GS is a mixture of signals from multiple brain regions without considering the possibility of spatial heterogeneity in the GS [9, 14, 16, 17, 18]. It has been shown that regressing out average-based GS results in negative correlations that do not have a biological basis and are artifacts in the voxels’ time series which lead to spurious findings [9, 16, 17]. In this paper, we showed that the AGSR method works voxel specifically and computes the neuronal correlations of the brain’s networks more accurately. This is because using the EMD method in computing AGS maximizes the spatial contributions to the GS. In other words, decomposing fMRI data in space using the FATEMD approach, which is done by considering features of each voxel’s neighbours, makes the computed AGS sensitive to brain regions’ heterogeneity.

When assessing transitivity and efficiency for different spatiotemporal domains of the fMRI data, no large differences in different temporal IMFs at the same spatial IMF were obtained. Thus, we concluded that the variability of transitivity and efficiency were just related to the spatial frequency domains. The high values of the transitivity and efficiency in the low spatial frequencies demonstrated the existence of the GS. On the other hand, high spatial frequencies, SIMF1 and SIMF2, represented the most network-specific data. Accordingly, the low spatial frequencies, SIMF3 to SIMF5 including all TIMFs, were considered as the AGS. Moreover, the observed averaged transitivity and efficiency values in the high spatial frequency domains were almost the same as the transitivity and efficiency values when GS is not removed from the fMRI time series. Considering these result, the summation of the low spatial frequency domains considered as the GS in the fMRI data just causes incorrect correlation between regions in the brain’s network and prevents reflecting the intrinsic property of the spatiotemporal nature of the fMRI data.

We examined the efficacy of our method by computing the seed-based functional connectivity for the TPN and TNN regions. Our results in agreement with previous studies [9, 53, 54], show that the negative correlations are intrinsic to the brain and do not appear just as a result of the GSR. We found that the AGSR method identifies the connectivity between the TPN and TNN regions according with the expected results of prior studies [9, 49]. We compared the connectivity results of the AGSR with the SGSR and when there is NR in the fMRI data. Despite the connectivity results of the SGSR method and when there is NR, applying our proposed method resulted in an enhancement to the detection of network-specific fluctuations of the brain. Furthermore, although the strength of the correlations is related to cognitive function, it has been shown [55] that the activity of the visual regions with the eyes-open rest condition is larger than with the eyes-closed rest condition. Our results showed high activation in visual regions in respect to the results of the SGSR and NR which were close to zero activity. In contrast, in auditory regions, lower activity seen in the result of applying AGSR appears to be related to the better removal of the acoustic noise heard by subjects during fMRI. This shows that the acoustic noise of the fMRI device which is almost constant in all TR times and interferes with auditory system activity can be removed through AGSR [56, 57]. Thus, AGSR method is able to correctly remove physiological and remained systemic noises after preprocessing without introducing artifactual correlations as confirmed by correlations between PCC and the reference regions.

In conclusion, AGS is a unique local signal for each voxel’s BOLD signal. In the AGSR method, the first and second spatial IMFs of each fMRI data, decomposed by FATEMD method, are simply summed up to have an fMRI data without GS. AGSR is a reliable method that works voxel-specifically for all subjects which leads to extract correct connectivity maps and provide accurate information about brain function.

There are some limitations to this Study, and alternative recommended strategies for future work that should be noted. In AGSR, the FATEMD and the ICEEMDAN approaches are applied to decompose the fMRI time series into their components to define the AGS. The FATEMD and the ICEEMDAN approaches are trivariate and univariate algorithms, respectively, while there is a multivariate EMD which is the extent of the EMD, applied for multivariate signal processing and can be utilized to decompose and process fMRI signals multivariately. Thus, instead of applying the FATEMD to compute SIMFS and then the ICEEMDAN method to compute TIMFs, it is possible to apply the MEMD to each fMRI image and compute spatiotemporal IMFs (STIMF). Although the multivariate analysis does not have volume nature and decompose the volume into one-dimensional signals [19], the sensitivity is increased by combining all data. MEMD by revealing possible inter-dependencies between signals provides useful information about the structure of their underlying system and helps to understand the complex organization of the brain.

Furthermore, in computing functional connectivity, we computed the Pearson’s correlation due to its simplicity and popularity. It also allows our findings to be comparable with other papers to test the efficacy of the proposed method. The Pearson’s correlation is used to compute the correlation coefficient between different regions to investigate how different brain regions are correlated with each other. It should be mentioned that the Pearson’s correlation assesses the strength of the linear relationship between two variables and is not able to indicate possible nonlinear relationships. In future work, we can use other correlation approaches that are sensitive to nonlinear relationships and are able to provide more detailed information. Besides, correlation computation does not imply causation. Causal relation computations can be used to investigate if a region or regions that are activated are caused by activation of other regions, because it is possible that some regions of the brain are activated together and correlate with each other, but does not mean one caused the others’ activation. Therefore, causality measurements by providing more information about how different brain regions are related to each other increase our knowledge about how the brain works and information flows in the brain networks. Thus, future work is needed to clarify the influence of other approaches.

Lastly, although the FATEMD and ICEEMDAN are optimized approaches for finding the best IMF sets, they still need more improvement in the sifting procedure to reduce the time of the calculation and to yield better decomposition performance. For instance, finding the optimum values of added white noise and the ensemble number to overcome the mode mixing problem and speed up the calculation in ICEEMDAN approach are two drawbacks of this approach. Therefore, the proposed method provides the opportunity to characterize the whole brain function. Future studies can be devoted to the application of our proposed method to the other image processing areas.

## 5. Acknowledgements

We are grateful to Doug Phillips for generous assistance in computational work on Compute Canada cluster and Jordan Chad for proofreading the manuscript. This work was partially supported by grant RGPIN-2015-05966 from Natural Sciences and Engineering Research Council of Canada. Data were provided [in part] by the Human Connectome Project, WU-Minn Consortium (Principal Investigators: David Van Essen and Kamil Ugurbil; 1U54MH091657) funded by the 16 NIH Institutes and Centers that support the NIH Blueprint for Neuroscience Research; and by the McDonnell Center for Systems Neuroscience at Washington University. The computational work was enabled by support provided of WestGrid (www.westgrid.ca) through Compute Canada (www.computecanada.ca).

